# Phenotypic complexity makes the molecular basis of evolutionary convergence predictable

**DOI:** 10.64898/2026.03.04.709483

**Authors:** Daohan Jiang, Matt Pennell, Lauren Sallan

## Abstract

Whether the outcomes of evolution are predictable remains a central question in evolutionary biology. Convergent evolution, in which similar phenotypes arise independently in distinct lineages, is often interpreted as evidence of predictability under shared selective pressures. However, empirical studies have shown that convergent phenotypes can exhibit similar or disparate molecular bases, and no general framework predicts when convergence should occur. Here, we show that phenotypic complexity determines the extent of convergence under shared conditions. Using a deliberately minimal abstraction linking genotype and phenotype and evolutionary simulations, we investigated how regulatory gene expression programs evolve under selection for shared phenotypic optima. We find that phenotypes of low complexity can be produced by diverse regulator expression programs, permitting phenotypic convergence despite molecular divergence. However, as the complexity of the target phenotype increases, the variety of viable regulatory solutions decreases, funneling independently evolving lineages towards similar expression programs. In this regime, molecular convergence emerges as a predictable consequence of selection acting on finite regulatory repertoires, consistent with the phenomenon of “deep homology” observed in structures like eyes and limbs. Allowing evolutionary changes in regulator-target interactions relaxes these constraints and enhances divergence in expression between phenotypically convergent species. Together, our results provide general expectations for predicting when phenotypic convergence should be accompanied by convergent genetic mechanisms, and shed light on the causes of convergence in a broad range of phenotypic traits.

## Introduction

A foundational problem in evolutionary biology is whether outcomes of evolution are generally predictable, especially over macroevolutionary timescales. The problem is encapsulated by the thought experiment of “replaying the tape of life”, which asks whether the same life forms would reappear if the evolution of life on Earth could start over [1]. One view, shared by Gould, emphasizes the role of historical contingency—that the outcome of evolution depends on historical events unrelated to present environments—and the resulting unpredictability, whereas an alternative view emphasizes the power of natural selection and the predictability of adaptive evolution under the same conditions [2, 3].

Many instances of convergent evolution (convergence), the phenomenon in which similar phenotypic states evolve independently in different lineages [4–6], appear to support predictability. In principle, convergent phenotypes must arise through mutations and be mediated through developmental processes and gene regulation. Yet, the relationships among genetic changes, phenotypic similarities, and the regime of selection remain unclear. Similar phenotypes may evolve via distinct genetic mechanisms, and the accessibility of particular genetic solutions may depend on the unique evolutionary history of a specific lineage [3, 7]. Furthermore, while phenotypic convergence is routinely interpreted as a result of adaptation to similar environments [5], it is unclear if a common regime of selection is sufficient or necessary for convergence to take place—does the same selective regime consistently favor the same solution, and could apparently similar phenotypes evolve under different selective regimes? Taken together, even when phenotypes appear “convergent”, the underlying evolutionary causes may reflect historical contingency rather than predictable adaptive responses. What is lacking is a general framework that connects evolutionary processes at the molecular and organismal levels and specifies when “true convergence” should be expected to occur.

Empirical work aiming to link evolutionary convergence at molecular and phenotypic levels has yielded mixed results. For example, comparative genomic and transcriptomic analyses of species with common ecological adaptations, such as echolocating mammals [8], marine mammals [9], and monogamous vertebrates [10], identified substitutions at the same loci toward the same sequence state and concordant expression level changes of the same gene in different species, which have been described as genomic or transcriptomic convergence. Subsequent studies, however, found little signal of genome- or transcriptome-scale convergence in these cases, once background expectations are accounted for [11–14]. The absence of broad signals has multiple interpretations that are not mutually exclusive: a lack of a shared genetic basis for independent adaptations, the involvement of too few genes or loci, or “noise” resulting from selection on traits other than the single adaptive trait under consideration [7, 15]. Detailed case studies found that the same loci, genes, or regulatory pathways are involved in producing other independently evolved phenotypes in closely related species or populations (e.g., [16–18], in which cases the convergence may also be referred to as parallel evolution or parallelism). This has also been found in a few cases involving moderately divergent species that have been separated for millions of years (e.g. [19–22]). Yet, in general, convergent molecular evolution is less likely between more distantly related lineages [18, 23–25]. Furthermore, studies examining genetic architectures of polygenic traits have shown that there can be many different genomic loci or genetic variants with similar effects on the same phenotypic trait (e.g., [26, 27]), indicating that the same phenotypic change can readily be produced via many genetic paths. Together, these studies show that convergence at phenotypic and molecular levels can be linked, but only to a limited degree and under a limited range of conditions, with no consistent expectations.

In light of these findings, the cases of evolutionary convergence known as “deep homology” appear particularly striking: homologous genes or gene families are involved in the development of functionally analogous, yet anatomically distinct structures that evolved independently in deeply divergent lineages [28, 29]. Some classic examples of deep homology include the involvement of shared developmental regulators be-tween vertebrate and arthropod appendages [30], and the sharing of developmental regulators between light-sensing organs (eyes) in diverse animal groups (vertebrates, cephalopods, arthropods, and cnidarians) [31– 33]. While the concept of deep homology was initially coined in the context of animal evolution, similar phenomena have also been documented in other groups, including plants [34] and fungi [35]. Deep homology is often explained by invoking homologous precursor structures that were present in a common ancestor of the species in question and served as building blocks for independently-evolved complex structures [29]. Under this view, the observed association between functional and molecular convergence reflects modification of pre-existing shared complexity rather than independent emergence under shared selective constraints. Some biological interpretations for the nature of such homologous precursors include cell types [29, 32] and modules in gene regulatory networks [36–39]. However, it is generally challenging to identify the hypothesized homologous precursors, as it is unclear where one should look for them. Furthermore, there is a possibility that the precursor is no longer present in its original form in any extant species. These challenges, together, limit the explanatory power of precursor-based hypotheses and leave unresolved whether deep homology must reflect historical contingency or can result from broader constraints on ways of achieving complex adaptation.

In this study, we show that deep homology, along with broader patterns of evolutionary convergence, can be explained by the interplay between fundamental properties of genotype-phenotype maps and natural selection. Using a model with deliberately minimal abstraction and making no specific assumptions about homologous precursor structures beyond shared regulatory genes, we explored how the expression program of regulatory genes would evolve in response to selection for a novel phenotype. We demonstrate that the most complex adaptations, perhaps counterintuitively, tend to be produced by similar gene expression programs in independently evolving lineages as a result of the limited capacity of the repertoire of regulatory genes to produce complex phenotypes. We also show that such a constraint on diversification can be reduced by evolutionary changes in regulator-target interactions. Taken together, our results unify disparate observations into general expectations for when convergence at different scales of biological organization should or should not arise under common selective pressures.

## Results

### A non-monotonic relationship between parallelism and phenotypic complexity

We considered how a novel character (a new body part, organ, or cell type) would evolve towards an optimal phenotypic state via evolutionary changes in local expression of regulatory genes (“regulators” hereafter) encoding *trans*-regulatory factors. Specifically, we asked whether independently evolving species will evolve similar or dissimilar regulator expression programs in response to selection for the same optimal phenotype. We started from simple scenarios where adaptation is mediated exclusively by evolutionary changes in local expression levels of the regulators as other potential genetic changes are too pleiotropic (thereby deleterious) to be acceptable. This would be the case if 1) the regulators are expressed in many other parts of the organism such that mutations in the regulators themselves are highly pleiotropic and deleterious [40] and 2) *cis*-elements interacting with these regulators are also pleiotropic and under similar constraints [41, 42]. Given these conditions, the optimum can only be achieved upon the emergence of a combination of expression levels of the regulators (“regulator expression program” hereafter) that produces an adaptive phenotypic state. Under such a setting, the regulator expression program (denoted as a vector **X** of length *n*, where each element represents the expression level of one regulator) that produces a given optimal phenotypic state (denoted as a vector **z**_*opt*_) can be obtained by solving a linear system **CX** = **z**_*opt*_. The matrix **C** characterizes effects of the regulators on the phenotype, with *C*_*i,j*_ representing the effect of the *j*-th regulator on the *i*-th phenotypic dimension (“trait” hereafter) per unit expression. Whether a solution exists and the dimensionality of the solution space when the system is indeed solvable will depend on the complexity of the selected phenotype (as measured by dimensionality of **z**_*opt*_ [43]). When **z**_*opt*_ has fewer dimensions than **X**, the system can have infinitely many solutions and the evolution of **X** can follow diverse paths (Fig. 1A). As dimensionality of **z**_*opt*_ increases, the solution space, in general, will have fewer dimensions; when **z**_*opt*_ has the same number of dimensions as **X**, there is at most a single solution, such that independently evolving lineages are expected to approach the same **X** eventually (Fig. 1B).

**Figure 1:**
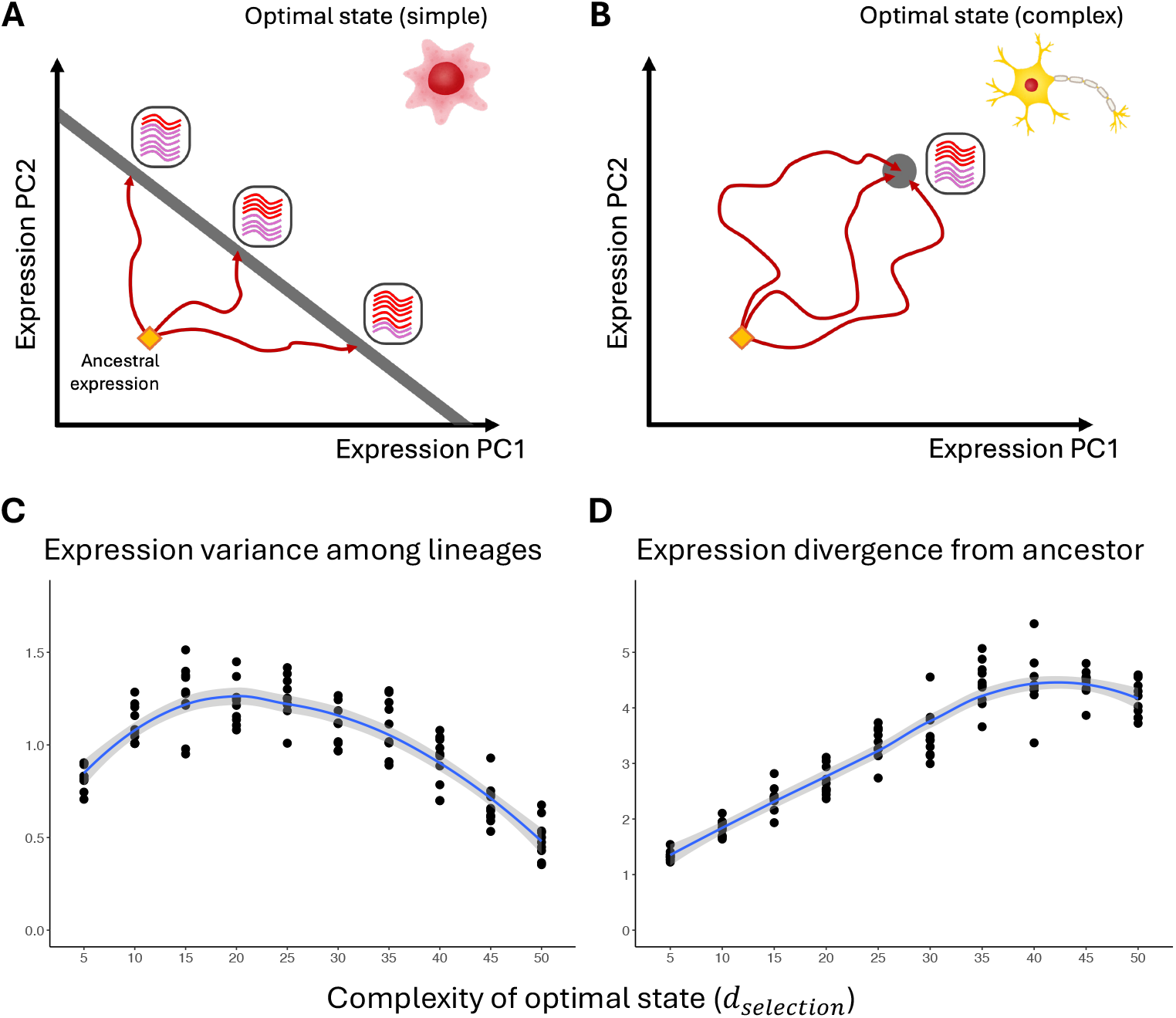
(A-B) Schematic illustrations for the evolution of novel regulator expression programs in response to selection on the phenotypic state. Axes represent two principal components of regulator expression. Red curves with arrows represent trajectories of evolution in different lineages. (A) The optimal state can be produced by various regulator expression programs, which fall along the gray line. Independent lineages (species) that evolved under the same regime of selection ended up at different points along the line, resulting in variation in regulator expression across lineages. (B) The optimal state can only be produced by a specific regulator expression program, which is represented by the gray dot. Different species evolved towards the only solution and there is little variation in expression programs among them. (C-D) Relationship between evolution of the regulator expression program and complexity of the optimal phenotypic state (quantified by the number of traits subject to selection, *d*_*selection*_). (C) Variance in regulator expression across species at the end of the simulation, plotted against *d*_*selection*_. (D) Divergence from the ancestral expression program at the end of the simulation, averaged across species and plotted against *d*_*selection*_. Each data point in (C) and (D) corresponds to a simulated scenario where a unique set of traits were under selection. Each curve in (C) and (D) is a locally estimated scatterplot smoothing (LOESS) curve, with the gray area representing the 95% confidence interval (CI). Cell images representing optimal phenotypes in (A) and (B) are from Irasutoya (https://www.irasutoya.com/.)

To confirm the above general predictions, we conducted evolutionary simulations with a total of 50 available regulators. We considered regimes of selection where the number of traits subject to selection (*d*_*selection*_) varied, and for each regime of selection, we simulated 20 replicate lineages (species) with the same ancestral expression program **X**_*a*_. At the end of the simulations, we examined variances of regulator expression levels among species as well as the mean level of expression divergence from the ancestral expression program **X**_*a*_. Our simulations revealed a non-monotonic relationship between expression variance and *d*_*selection*_, where expression variance first increased, peaking at an intermediate *d*_*selection*_, and then decreased (Fig. 1C). The degree of expression divergence from the ancestor mostly increased in the range of parameters examined, only dipping when more than 40 traits were selected (Fig. 1D). Together, there is a range of *d*_*selection*_ in which the regulator expression program evolved farther away from the ancestral one but closer to each other as *d*_*selection*_ increases. The mean fitness across replicate lineages 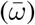 at the end generally decreases with *d*_*selection*_, especially when *d*_*selection*_ exceeds 20 (Fig. S1), demonstrating a cost of complexity [43] and stronger constraint on the speed of adaptation at higher *d*_*selection*_.

### Phenotypic complexity drives convergence from diverse ancestral states

The above simulations assumed that evolution in different species started from the same ancestral state, which, in general, is expected to make the evolved species more similar to each other [44]. To investigate whether lineages evolving from divergent ancestral states show similar patterns of convergence, we performed two-stage evolutionary simulations: in stage 1, regulator expression programs in different species undergo independent neutral evolution and reach high expression variance; in stage 2, selection on the phenotypic state is imposed. Divergence in stage 1, while referred to as neutral for simplicity, can also be interpreted as divergence due to different regimes of selection during stage 1 and/or that the novel character is derived from non-homologous developmental progenitors with different regulator expression programs to start with.

We expected that, when *d*_*selection*_ is small and there are many expression programs that can produce the optimal phenotype, different species will evolve towards expression programs that are relatively close to their respective expression programs at the end of stage 1, thereby preserving expression variance among species (Fig. 2A); in contrast, when *d*_*selection*_ is high such that there is a single optimal expression program, different species will eventually converge towards this optimum regardless of their prior divergence (Fig. 2B). Together, we predicted that the expression variance across species at the end will generally decrease with *d*_*selection*_, which is confirmed by our simulations (Fig. 2C). Expression divergence of each species from its own ancestral state, on the other hand, generally increased with *d*_*selection*_ (Fig. 2D), which means each species underwent more evolutionary change in regulator expression in response to selection for more complex optimal states. Together, the complexity of the selected phenotype leads not only to parallel expression evolution from a common ancestral state, but also to convergence from diverse ancestral states.

**Figure 2:**
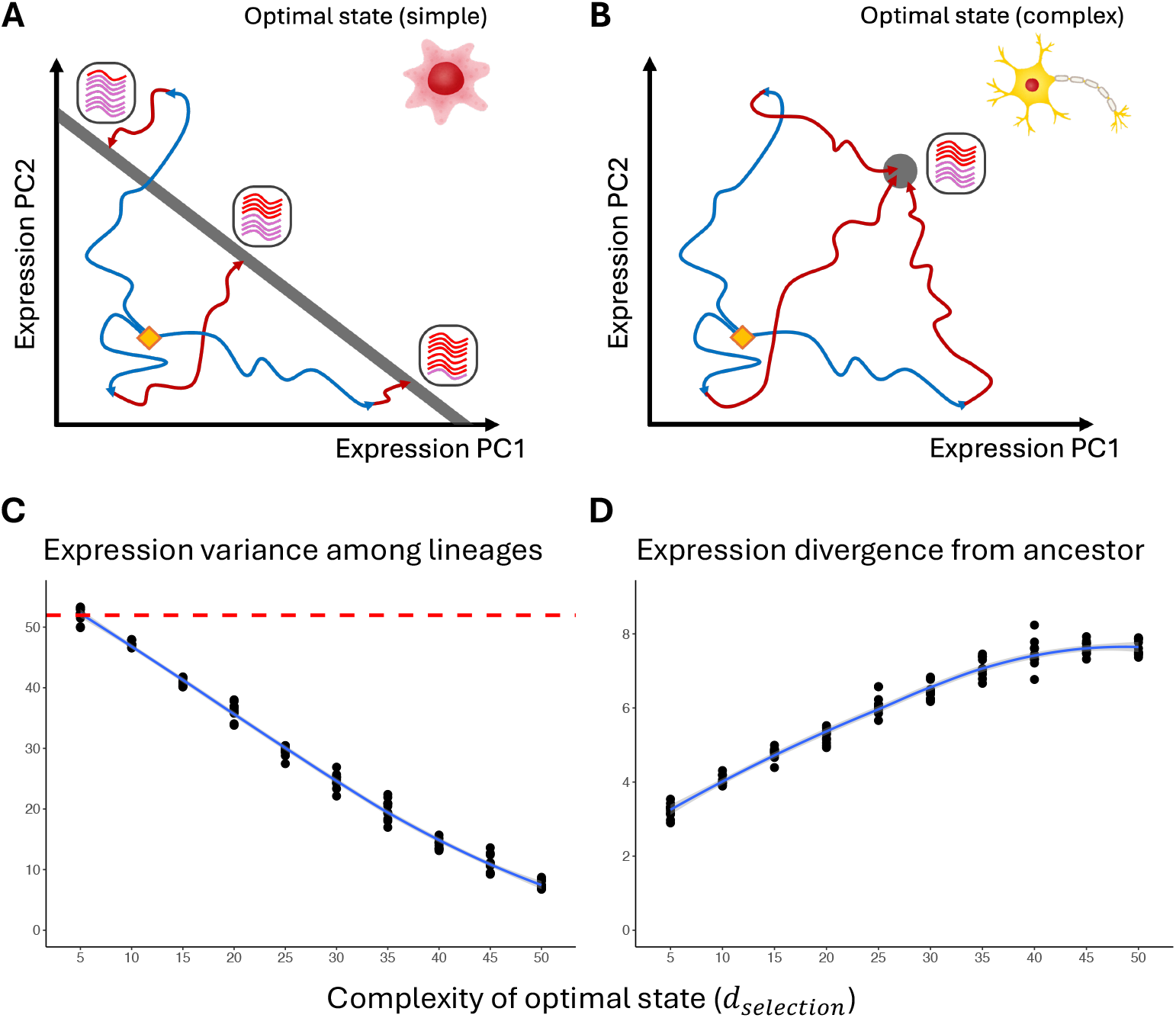
(A-B) Schematic illustrations for two-stage evolution of novel regulator expression programs. Blue curves with arrows represent trajectories of neutral evolution in stage 1 (before phenotypic traits of interest are subjected to selection), whereas red curves with arrows represent trajectories of evolution in stage 2 (after selection is imposed). (A) The optimal state can be produced by various regulator expression programs (gray line). (B) The optimal state can only be produced by a specific regulator expression program (gray dot). (C-D) Relationship between evolution of the regulator expression program and complexity of the optimal phenotypic state (*d*_*selection*_). Notations are the same as Fig. 1C-D, unless stated otherwise. (C) Variance in regulator expression across species at the end of stage 2, plotted against *d*_*selection*_. Red dashed line represents expression variance right before selection is imposed (at the end of stage 1 and the beginning of stage 2). (D) Divergence from the expression program at the end of stage 1, averaged across species and plotted against *d*_*selection*_.

### Evolution of phenotypic effects reduces expression convergence

Lastly, we examined how evolutionary changes in the phenotypic effects of the regulators would affect evolvability and the relationship between expression divergence and phenotypic complexity. To this end, we performed simulations where elements of the **C**-matrix were affected by mutations, which can be interpreted biologically as those in tissue-specific *cis*-elements. These evolutionary changes, even if rare, can lift the strict limit to phenotypic evolution imposed by the number of available regulators. While a non-monotonic relationship between expression variance among species and the complexity of the selected phenotype is recovered, the curve is dampened compared to that when **C** did not evolve, with higher expression variance at high *d*_*selection*_ (Fig. 3A-B), indicating that evolutionary changes in **C** did enhance expression divergence between species. Divergence from the ancestral expression program, on the other hand, showed a similar trend as that seen when **C** did not evolve (Fig. 3C-D). Allowing **C** to evolve is also expected to increase the overall input of potentially adaptive mutations and enable faster adaptation. Indeed, fitness at high *d*_*selection*_ ended up higher when **C** could evolve (Fig. S3).

**Figure 3:**
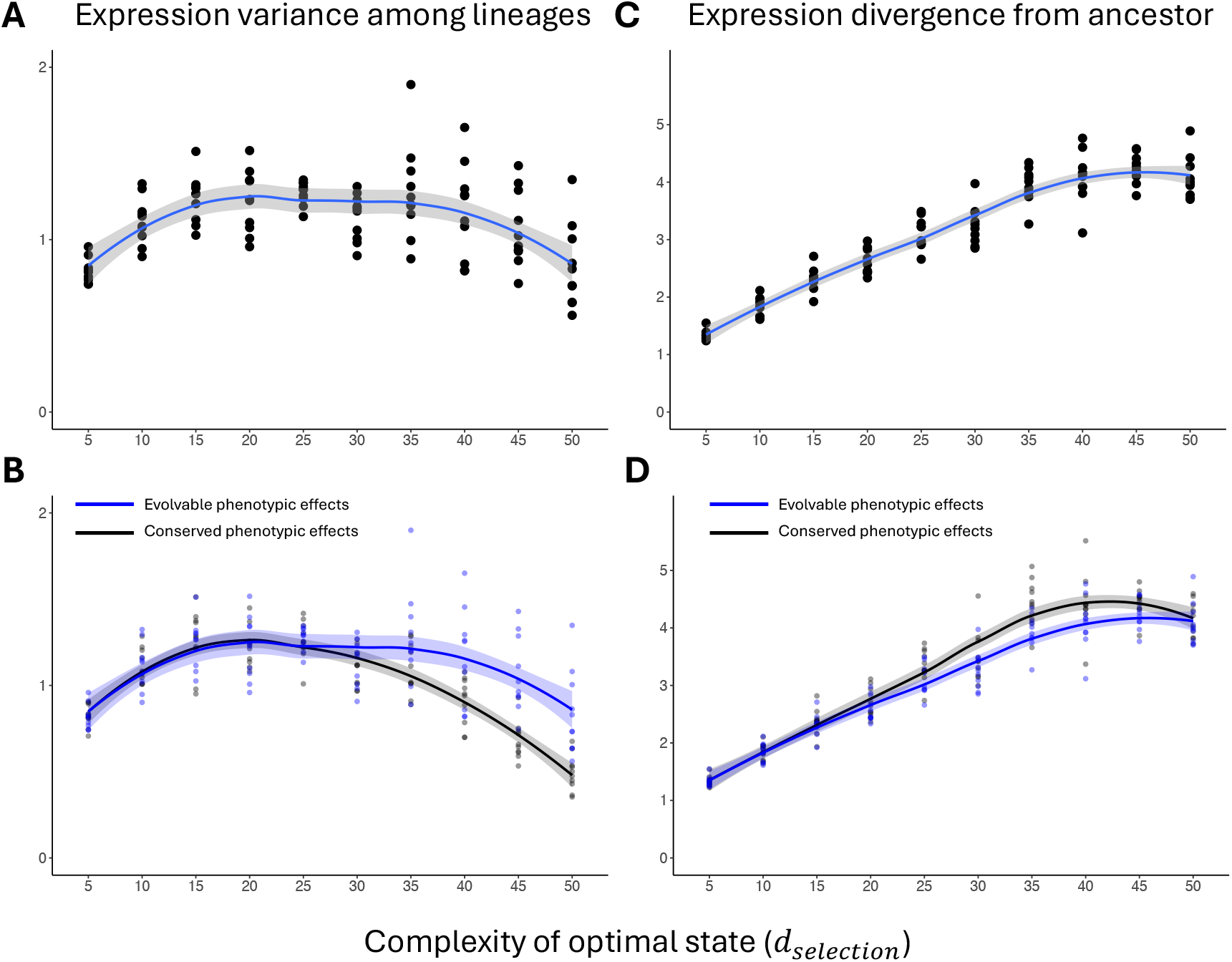
Effect of complexity on regulator expression evolution when phenotypic effects of the regulators (elements of the **C**-matrix) can evolve. (A) Variance in regulator expression across species at the end of the simulation, plotted against *d*_*selection*_. (B) Comparison of the curve in (A) (blue) with that from simulations with constant **C** (black curve and dots; the same as those in Fig. 1C). (C) Divergence from the ancestral expression program at the end of the simulation, averaged across species and plotted against *d*_*selection*_. (D) Comparison of the curve in (C) (blue) with that from simulations with constant **C** (black curve and dots; the same as those in Fig. 1D). CIs of LOESS curves in (B) and (D) are in colors that match the corresponding curves but lighter.

## Discussion

Our simulations recreated different types of observed evolutionary convergence, revealing minimal circumstances under which they can predictably occur (summarized in Table 1 and Fig. 4). Parallelism, in which species with a common ancestral state evolved similar adaptive phenotypes with similar underlying regulator expression programs, can happen when the selected phenotype is of either low or high levels of complexity (Fig. 1C). A second type, which we refer to as function-only convergence, is when species evolved toward the same pre-set adaptive phenotype but achieved it with different underlying regulator expression programs. This type of convergence happened when species started from the same ancestral state and the selected phenotype is of moderate complexity (Fig. 1C, middle), or when species with different ancestral states at the onset of selection evolved towards a shared optimal phenotype of low to moderate complexity (Fig. 2C, left side). The final type is deep homology: as complexity of the selected phenotype increases, the space of regulator expression programs that can produce it shrinks, causing species with different ancestral expression programs to converge at both phenotypic and regulatory levels (Fig. 2C, right side). It should be noted that these different types of convergence are not discrete categories but exist along a continuous spectrum, and there is no expression variance or ancestral divergence cutoff to delineate them (Fig. 4). However, to have a distinction between them may be useful for formulating future studies of the causes of any particular case of phenotypic convergence.

**Table 1:** Summary of different types of evolutionary convergence, and which cases in Fig. 1C and Fig. 2C they correspond to. “Not specified” means the concept does not have specific assumptions about the subject.

| Type | Same ancestral states? | Similar genetic mechanisms? | Similar phenotypes? | Complexity of shared phenotype | Representation in variance-complexity plots |
| --- | --- | --- | --- | --- | --- |
| Convergence (broad-sense) | Not specified | Not specified | Yes | Not specified | Fig. 1C and Fig. 2C |
| Parallelism | Yes | Yes | Yes | Low or high | Fig. 1C, left and right side |
| Function-only convergence | Not specified | No | Yes | Low or moderate | Fig. 2C, left side |
| Deep homology | No | Yes | Yes | High | Fig. 2C, right side |

**Figure 4:**
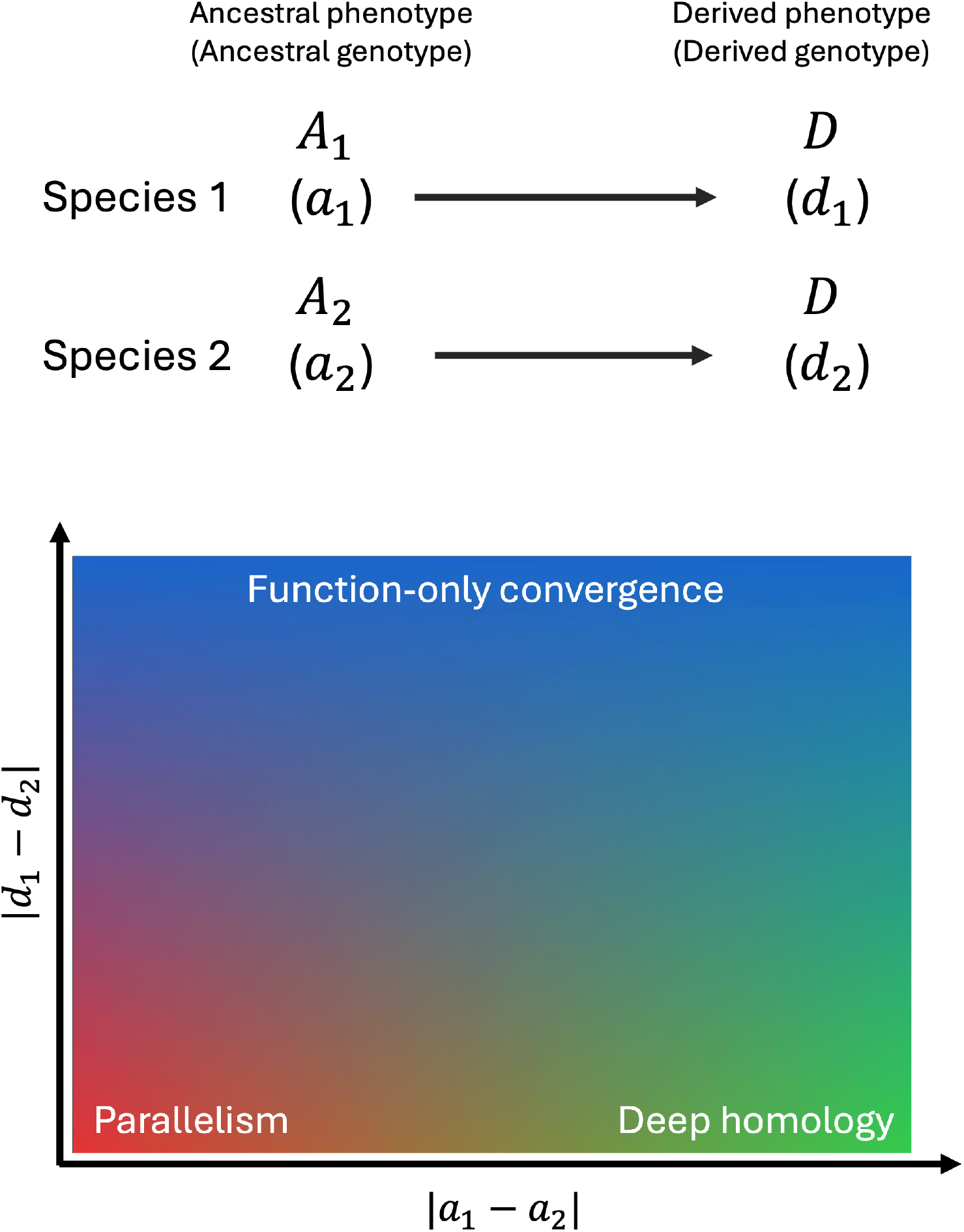
Schematic illustration of the distinction between different types of evolutionary convergence. Upper part: A scenario where two species with ancestral phenotypes *A*_1_ and *A*_2_ underwent convergent evolution toward the same derived phenotype *D*. Genotype underlying each phenotype is shown in parentheses. Lower part: Parameter space corresponding to different types of convergence. The X-axis represents the difference between ancestral genotypes ( |*a*_1_−*a*_2_| ), and Y-axis represents the degree of difference between derived genotypes (| *d*_1_−*d*_2_| ). When |*a*_1_−*a*_2_| and |*d*_1_−*d*_2_| are both small, the case of convergence is considered parallelism. When |*a*_1_−*a*_2_ |is high but |*d*_1_−*d*_2_ |is small, it is deep homology. When |*d*_1_−*d*_2_| is large, it is function-only convergence.

Of the three types of convergence, function-only convergence and deep homology often result in similar but structurally distinct phenotypes, unlike in our simulation where species evolved toward a single adaptive phenotype. Given our interest in broad-scale convergence, the adaptive phenotype in our simulation can and should be interpreted as representing a “core function” that is subject to selection—e.g., it better represents the function of imaging than eye anatomy as a whole. Once a core functional trait evolves, additional, supporting structures commonly perceived as belonging to the same body part can evolve under “internal selection” imposed by the focal trait [45]. These can then take on different overall forms in different species because of different ancestral conditions of divergent lineages, layering contingency atop predictability. For example, the sclerotic ring that supports vertebrate eye requires internal ossification, such that arthropods and mollusks cannot produce the same supporting structure even under the same selective pressure [46]. Furthermore, even traits related to the core function can vary among species that exhibit convergence due to differences in the regimes of selection. This is likely to occur since environments where these species evolved were likely not exactly the same. In the case of eyes, while light sensing and later enhancements in visual acuity have been selected for in multiple groups of animals, selection for specific aspects of vision such as sensing of different wavelengths and polarized light have varied across lineages [47]. To tease apart the effects of these factors in specific cases of convergence can be challenging, as it is often unclear which aspects of the measured phenotype are relevant to the functions of interest. It would require detailed investigations of the fitness consequences of different aspects of the phenotype.

While the adaptive and molecular drivers of phenotypic convergence were set for the simulated species, it may be difficult to determine the same for real species or provide an exact categorization of their convergence. This is because the ancestral states of the species of interest, the adaptive significance of their similar phenotypes, and the genetic mechanisms underlying these phenotypes are often unclear. This means one cannot describe a species as exhibiting a specific type of convergence without targeted research into both the molecular underpinnings and past states of the target trait. While our results should be useful for most recognized cases of apparent convergence, there are instances where phenotypic similarity can result from distinct functional demands. For instance, limbless forms have evolved multiple times in tetrapods, including in squamates (snakes and multiple groups of limbless lizards) [48], amphibians (caecilians and limbless salamanders) [49], and stem tetrapods like aistopods [50]. Limbless forms have been proposed to be adaptations to either swimming or burrowing, as both lifestyles are observed in limbless lineages and there is mixed fossil evidence; this suggests that conditions in both settings can favor the same phenotypic solution, and the outcome of evolution in one setting can be readily “repurposed” for the other. It also implies that, for specific groups of interest, it can be challenging to confirm which of several environments or sets of functional demands was the primary driver of limb loss, which relies on the availability as well as interpretation of fossil records (e.g., snakes [51, 52]). Genetic mechanisms of limb loss are also unknown for most of these lineages, with the exception of snakes [53], further limiting categorization. Furthermore, convergence, in principle, can happen by chance in the absence of selection, which has been shown to be prevalent in the evolution of protein sequences as well as morphological characters with a small number of potential states [11, 12, 54]; such convergence does not fall into any of the scenarios presented in this study. Together, a broadly defined concept of convergence will still be useful for describing observations of convergence, especially when ancestral states of the species under concern or adaptive significance of their phenotypes is unknown. We also suggest that, when it is unclear whether a similarity between species represents convergence or homology (e.g., the shared state had a single origin but was lost in some lineages [55], or there was incomplete lineage sorting [56]), the above terms for convergence should not be applied; rather, the observation should only be described as a homoplasy before the evolutionary history is confirmed.

Deep homology, as discussed above, entails the appearance of similar regulatory expression programs underpinning functionally similar body parts or cell types in divergent species, despite the absence of comparable phenotypes in their common ancestor. Such similarity in gene expression programs can also evolve between different body parts within the same organism as regulators expressed in pre-existing body parts are co-opted for a novel structure or function (e.g., origin of the eye spots on butterfly wings [57] and the posterior lobe in fruit flies [58]). Although this phenomenon has been discussed as a type of deep homology [59], it should be noted that convergence between species and the evolution of (dis)similarity between parts of the same organism are distinct aspects of evolution that may have different driving mechanisms. These two aspects of phenotypic evolution are both constrained by the available repertoire of regulators but take place under different regimes of selection. Evolutionary innovation within the same organism is commonly driven by selection on a function that is not performed by pre-existing body parts [60], as in the aforementioned examples. In this study, we focused on a single body part in each species, and mechanisms driving (dis)similarity between parts of the same organism should be the subject for future studies.

Our simulations showed that the number of distinct regulators relative to the complexity of phenotypic traits under selection is a limiting factor that constrains adaptation and drives convergence in expression pro-grams. However, if the same regulator evolved different phenotypic effects in different species, the space of regulator expression programs that produce a given adaptive phenotype would be different between species as well. As a result, there is less parallelism in regulator expression in complex adaptation (Fig. 3). In this study, we did not make specific assumptions about the mutational origins of these evolutionary changes. Phenotypic effects of the regulators can be modified by *cis*-regulatory mutations that alter the regulators’ effects on the expression of the effector genes responsible for producing the phenotype. The origin of tissue- or cell type-specific *cis*-elements is particularly relevant, as mutations in *cis*-elements that are already used in other tissues or cell types are more likely to have deleterious pleiotropic effects. There remains much to be discovered, however, about the dynamics of the birth of novel *cis*-elements, making it challenging to parameterize when modeling this evolutionary process. There has been recent progress made in predicting regulatory potential of putative *cis*-elements as well as detecting selection on regulatory activities [61–68], which may shed light on how frequently tissue-specific *cis*-elements evolve and their role in adaptation and evolutionary innovations. Another potential mechanism by which local effects of regulators could change is through change in allosteric regulation, which was likely involved in the evolutionary innovation of placenta in placental mammals [69]. As the number of available regulators is a major source of constraint on pathways to adaptation, expansion of the repertoire of regulators via gene or genome duplication is presumably an essential means by which the constraint can be lifted. It has been suggested that genome duplication enabled evolutionary innovation and diversification in certain groups (e.g., teleost fishes [70] and flowering plants [71]), yet the validity of the causal relationship and likely mechanisms by which genome duplication could enhance evolvability have been debated [72–74]. The effect of gene and duplication on the dynamics of evolutionary convergence is likely affected by multiple additional factors that would be challenging to parameterize, such as epistatic interactions between duplicates (e.g., redundancy) and evolutionary changes (e.g., loss of a copy and change in tissue specificity) that took place before the onset of selection for novel functions. Modeling these processes is beyond the scope of this study but will be of great interest to future research.

In this study, we investigated principles of convergent evolution in the origin of novel phenotypes under selection. Using a deliberately simplified modeling framework and evolutionary simulations, we demonstrate the conditions under which different types of convergence can take place. Our findings offer a general explanation and a constraint-mediated basis for the predictable occurrence of for the often puzzling phenomenon of deep homology. Our results show that, given constraints on regulatory solutions and a shared optimal phenotypes, convergence predictably occurs. This suggests that, even phenotypic evolution itself is predictable to a degree, whether the underlying genetic mechanisms are shared or divergent depends on the complexity of the adaptive solution and the starting conditions. These findings should inform the interpretation of the origins of convergent traits in different contexts and help distinguish the pathways leading to functionally identical solutions across the tree of life.

## Methods

### Model of gene regulation

Let there be *n* regulatory genes (regulators) that are not directly involved in producing the phenotype and *m* effector genes (effectors) regulated by the regulators. Their expression levels are represented by two column vectors, 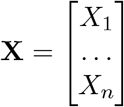 and 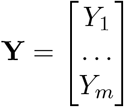, respectively. The vector **X** as a whole specifies the identity of a body part or cell type, and is expected to be the same among homologous body parts or cell types in different species, and to be different between phenotypically distinct parts in the same organism [59, 75–78].

Per-unit-expression effect of the regulators on the expression of the effectors is captured by a matrix **A**, where element *A*_*i,j*_ is the *j*-th regulator’s effect on the *i*-th effector’s expression. Expression dynamics of the effectors can be described by a system of ordinary differential equations, following previous studies [79, 80]. For the *i*-th effector, the rate of change of its gene product (RNA or protein) abundance is given by

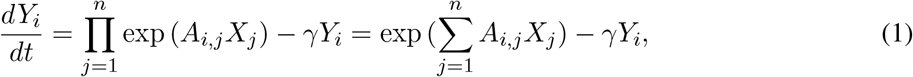

where *γ* is the decay rate of the gene product. The term exp (*A*_*i,j*_*X*_*j*_) represents the regulatory effect of the *j*-th regulator on the *i*-th effector, with *A*_*i,j*_ representing regulatory effect per unit expression. When *A*_*i,j*_ > 0, exp (*A*_*i,j*_*X*_*j*_) increases with *X*_*j*_ and the regulatory effect is activation; when *A*_*i,j*_ < 0, exp (*A*_*i,j*_*X*_*j*_) declines with *X*_*j*_ and the regulatory effect is repression. When exp (*A*_*i,j*_*X*_*j*_) = 1, which is reached when *X*_*j*_ = 0 or *A*_*i,j*_ = 0, the *j*-th regulator has no regulatory effect on *Y*_*i*_. The equilibrium of *Y*_*i*_, which is reached when 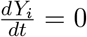, is

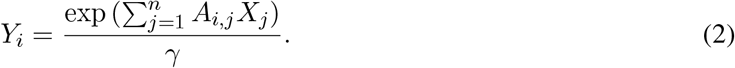

When *γ* = 1, as assumed throughout this study, the above equation becomes

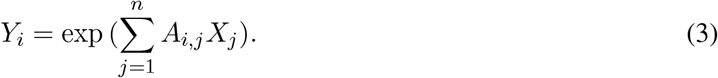

Together, there is

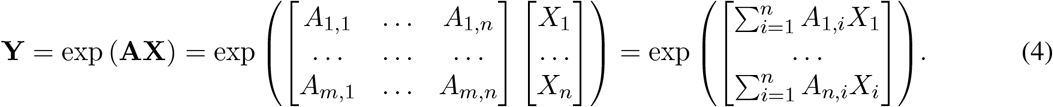

Here, **A** is an *m*× *n* matrix that summarizes the strength and direction of each regulator’s effect on each effector.

The **A**-matrix can be further decomposed into the effect of *cis*-elements on effector expression and strength of interaction between *cis*-elements and *trans*-factors encoded by the regulators. Let there be *L cis*-elements that can potentially regulate the expression of the *i*-th effector. The regulatory parameter *A*_*i,j*_ can be written as

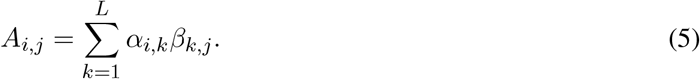

Here, *α*_*i,k*_ captures the effect of the *k*-th *cis*-element on the *i*-th effector (per unit expression of the *j*-th regulator), with its sign capturing whether the effect is activation or repression and its absolute value capturing the magnitude of the regulatory effect. Strength of interaction (binding affinity) between the *k*-th *cis*-element and the *j*-th regulator is captured by *β*_*k,j*_ ≥ 0. Together, there is

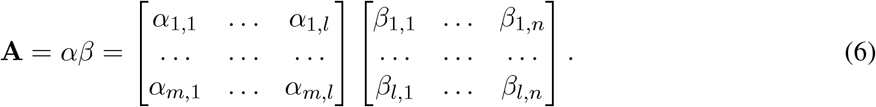

The observed phenotypic state is modeled as a linear combination of log-transformed expression levels of the effectors. For a given phenotypic trait *z*_*i*_, there is

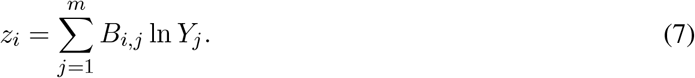

Here, *B*_*i,j*_ is a parameter characterizing the per-unit-expression effect of the *j*-th effector on *z*_*i*_. When *B*_*i,j*_ > 0, *z*_*i*_ is a monotonically increasing but concave function of *Y*_*j*_; when *B*_*i,j*_ < 0, *z*_*i*_ is a monotonically decreasing but convex function of *Y*_*j*_. When *B*_*i,j*_ = 0, *Y*_*j*_ has no effect on *z*_*i*_. Together, the effect of *Y*_*j*_ on *z*_*i*_ is expected to have a diminishing-return property, which is widespread in genotype-phenotype maps [81, 82]. Note that *z*_*i*_ is meant to represent normalized trait values, but not quantities that have to be non-negative such as body mass or absolute abundance of a chemical.

If the phenotype of interest is *d*-dimensional, Eqn. (7) becomes

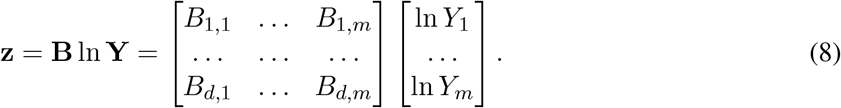

Here, **z** is a vector of length *d* and **B** is a *d*× *m* matrix that summarizes the per-unit-expression effects of the effectors on the trait values. Substituting **Y** with exp (**AX**), Eqn. (8) becomes

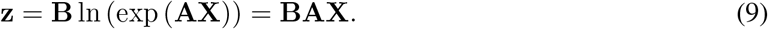

### Selection on the phenotype

Fitness is a multivariate Gaussian function of the character state **z**. The strength of selection along different phenotypic dimensions was characterized by a covariance matrix **S**. Fitness, denoted as *ω*, is calculated as

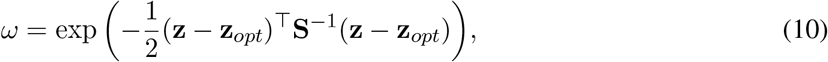

where the column vector **z**_*opt*_ is the optimal character state. Note that phenotypic dimensions that are not subject to selection will not be included in the above calculation.

Given associativity of matrix multiplication, Eqn. (9) can be written as **z** = (**BA**)**X**. With regulatory parameters (**A**) held constant, per-unit-expression phenotypic effect of the regulators can be summarized with a *d×n* matrix **C** = **BA** where each row corresponds to a phenotypic dimension (trait) and each column corresponds to a regulator. Assuming that the effects of mutations on each element of **X** have a continuous distribution such that any **X** ≥ 0, in principle, can be achieved via mutations. The optimal state, represented by a vector **z**_*opt*_, is achievable if the system **CX** = **z**_*opt*_ is solvable. While solvability of the system will depend on the structure of **C** and **z**_*opt*_, dimensionality of the solution space, in general, is expected to decrease with the dimensionality of **z**_*opt*_.

### Evolutionary simulations

Before simulations, we generated a **C** matrix that is 1000 × 50. The number of regulators is of the same magnitude as those of known developmental regulator gene families like the *Hox* family [40, 83]. We sampled 25 numbers (*C*_1_, …, *C*_25_) from an exponential distribution Exp(2) and then randomly shuffled a vector containing *C*_1_, …, *C*_25_ and −*C*_1_, …, −*C*_25_ to obtain 1000 permutations, each becoming a row of **C**.

All simulations started with an ancestral regulator expression vector **X**_*a*_ = **1**, unless stated otherwise. When the number of phenotypic dimensions subject to selection is *d*_*selection*_ (**z**_*opt*_ is *d*_*selection*_-dimensional), *d*_*selection*_ traits were randomly drawn from the 1000 traits. Elements of **z**_*opt*_ that correspond to traits under selection are equal to 2. With all elements of **z**_*opt*_ being positive, **CX** = **z**_*opt*_ will not be rendered unsolvable due to sign compatibility and its solvability will only depend on the dimensionality of **z**_*opt*_ relative to **X**. For each regime of selection, 20 replicate lineages (species) were simulated. For each *d*_*selection*_, we drew 10 sets of traits to serve as traits under selection, and simulated evolution with each set of traits under selection to account for random fluctuations in mutational covariance between traits under selection.

Simulations were performed following a sequential fixation model [84]. The total number of mutations that would occur during each simulation was drawn from a Poisson distribution with mean equal to 2*T N*_*e*_*nµ*, where *n* = 50 is the number of regulatory genes considered, *T* = 2×10^6^ is the duration of the simulation in terms of time steps, *N*_*e*_ = 1000 is the effective population size, and *µ* = 5×10^−6^ is the rate of mutations affecting regulator expression (per regulator per genome per time step). For simplicity, we assumed that each mutation can affect only one element of **X**, and all elements have the same probability of being affected. When a mutation affecting *X*_*i*_ occurs, a number *δ* would be sampled from a normal distribution 𝒩 (0, 0.1), and *X*_*i*_ would be multiplied by exp *δ*. The fitness of each genotype was calculated in the same way as Eqn. (10), with diagonal elements corresponding to traits under selection equal to 100 and off-diagonal elements equal to zero. Fixation probability of each mutation was calculated following Kimura’s method [85]:

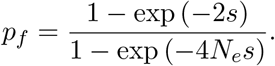

The coefficient of selection of the mutation, *s*, was calculated as 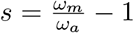, where *ω*_*m*_ and *ω*_*a*_ are mutant and ancestral fitness, respectively.

At the end of the simulations, we calculated cross-lineage variances of each element of ln **X**. The sum of these variances was used to represent the degree of expression variation among lineages. For each lineage, we computed the Euclidean distance between ln **X** and ln **X**_*a*_ to represent the degree of overall expression divergence from the ancestor [86, 87] and took the average across lineages for each regime of selection.

### Expression divergence before the onset of selection

In the two-stage simulations, evolutionary divergence of **X** during stage 1 was modeled as a multivariate Brownian motion process, and evolution in stage 2 was modeled in the same way as described in the previous section. The regulator expression programs at the end of stages 1 and 2 are denoted as **X**_1_ and **X**_2_, respectively. At the end of stage 1, which lasts for time *T*_0_, there is 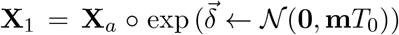, where **m** is a covariance matrix characterizing the rate of evolution and 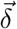 is a vector sampled from the corresponding multivariate normal distribution. In this study, we set *T*_0_ = 100, with all diagonal elements of **m** equal to 0.01, and all off-diagonal elements of **m** equal to zero. The expected total expression variance among lineages at the end of this episode of divergence is thus 50, which is close to what was obtained in our simulations (Fig. 2, red dashed line). At the end of the two-stage simulations, we calculated, for each lineage, the Euclidean distance between its ln **X**_2_ and ln **X**_1_ as a measurement for the amount of expression evolution under selection, and took the average of lineages for each regime of selection.

### Evolutionary changes in phenotypic effects of regulators

In our simulations where **C** co-evolved with **X**, we assumed that each mutation that affects **C** affects only one element. When a mutation affecting *C*_*i,j*_ occurs, a number *δ* would be sampled from a normal distribution𝒩 (0, 0.1), and *C*_*i,j*_ would become *C*_*i,j*_ + *δ*. The mutation rate was set as 10^−7^ per element of **C** per time step. Allowing elements of **C** to mutate independently is expected to mitigate constraint on evolution and reduce evolutionary convergence in **X**; thus, results obtained under such a setting will provide a conservative view of constraint and convergence.

All analysis of simulation results and generation of plots were done in R [88].

## Acknowledgment

We thank Günter Wagner, Joanna Wolfe, and Mark Kim for feedback on earlier versions of this manuscript. D.J. and L.S. are supported by Okinawa Institute of Science and Technology Graduate University.

## Data and code availability

Code and data files used in this study are available at https://github.com/RexJiangEvoBio/deep_homology.

## Supplementary Materials

**Figure S1:**
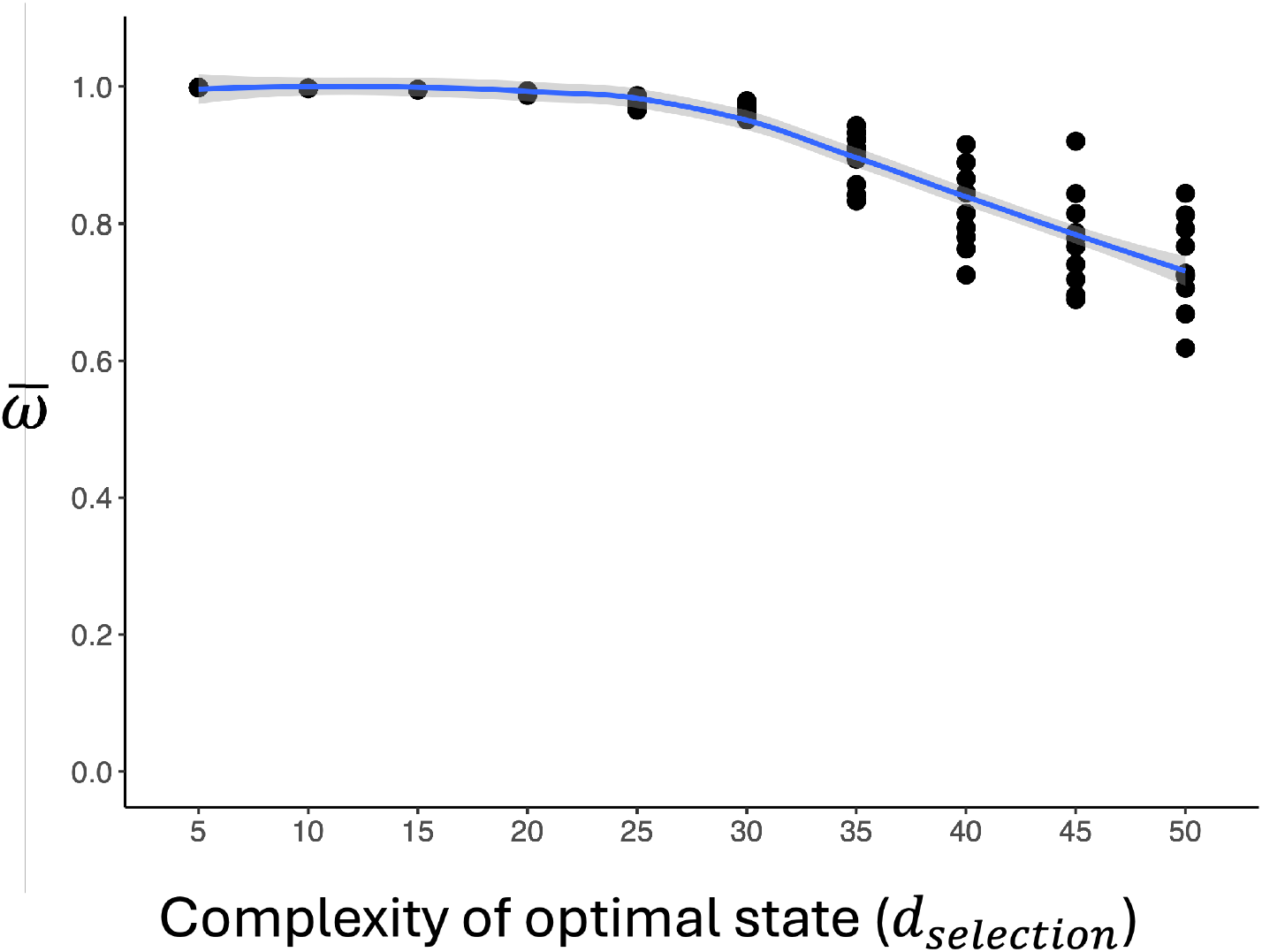
Mean fitness across species 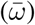at the end of simulations, plotted against *d*_*selection*_. Each data point corresponds to a simulated scenario where a unique set of traits were under selection. The curve is a locally estimated scatterplot smoothing (LOESS) curve, with the light gray area being the 95% confidence interval (CI).

**Figure S2:**
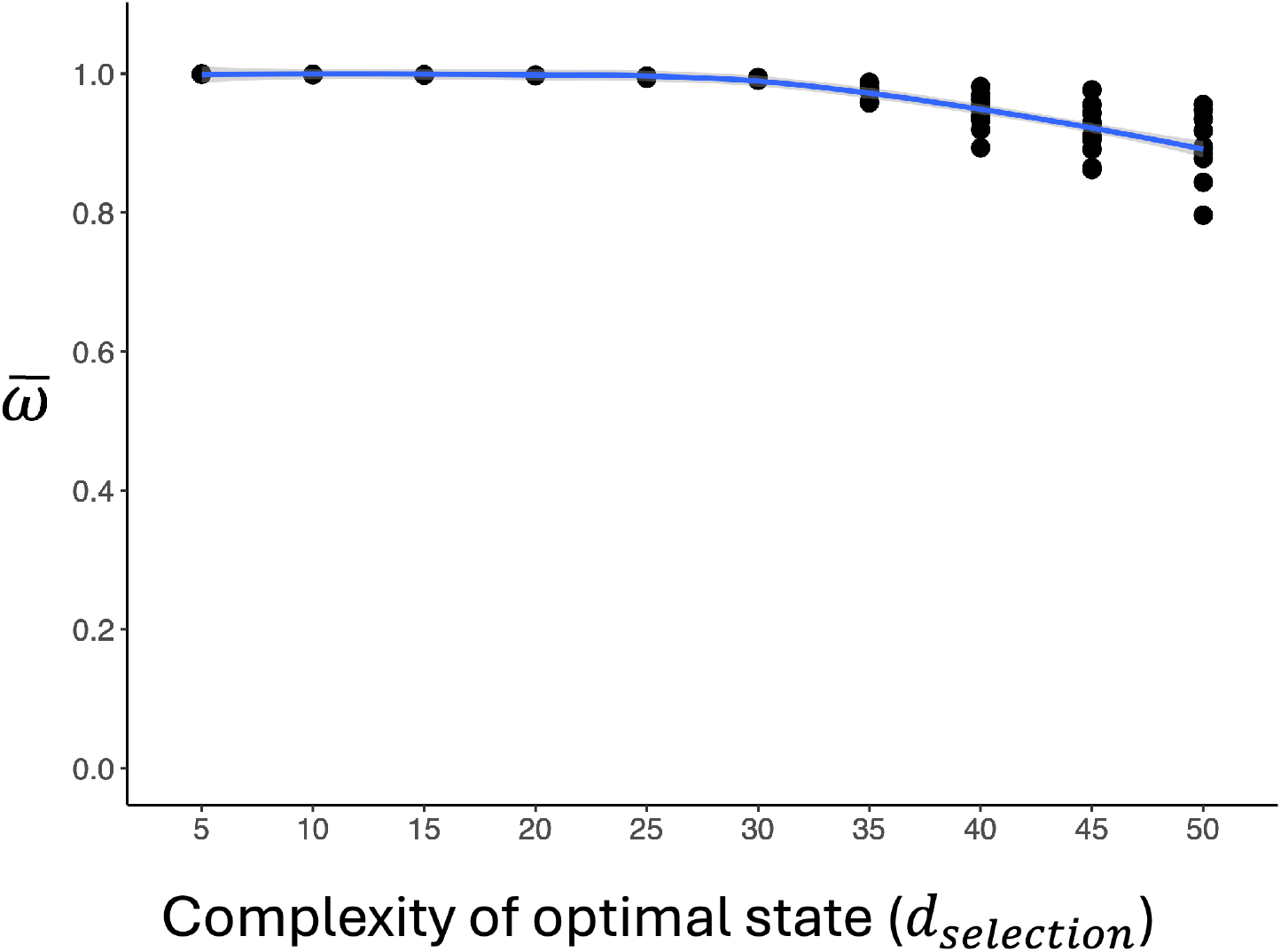
Mean fitness across species 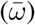 at the end of stage 2 in two-stage simulations, plotted against *d*_*selection*_. Each data point corresponds to a simulated scenario where a unique set of traits were under selection. The curve is a locally estimated scatterplot smoothing (LOESS) curve, with the light gray area being the 95% CI.

**Figure S3:**
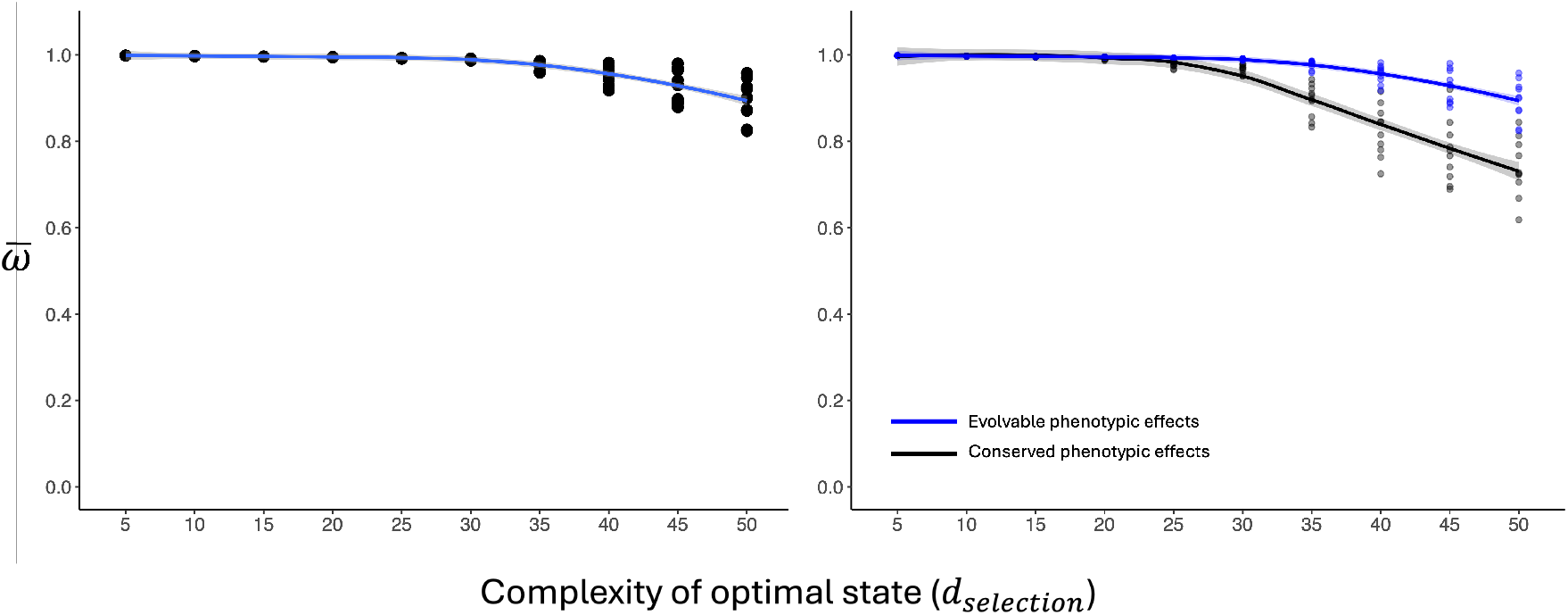
(A) Mean fitness across species 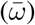 at the end of simulations with the **C**-matrix evolving, plotted against *d*_*selection*_. Each data point corresponds to a simulated scenario where a unique set of traits were under selection. The curve is a locally estimated scatterplot smoothing (LOESS) curve, with the light gray area being the 95% CI. (B) Comparison of the curve in (A) (blue) and that in Fig. S1.

